# Serum from COVID-19 patients early in the pandemic shows limited evidence of cross-neutralization against variants of concern

**DOI:** 10.1101/2021.11.10.468174

**Authors:** Amanda J. Griffin, Kyle L. O’Donnell, Kyle Shifflett, John-Paul Lavik, Patrick M. Russell, Michelle K. Zimmerman, Ryan F. Relich, Andrea Marzi

## Abstract

Severe acute respiratory syndrome coronavirus 2 (SARS-CoV-2) results in a variety of clinical symptoms ranging from no or mild to severe disease. Currently, there are multiple postulated mechanisms that may push a moderate to severe disease into a critical state. Human serum contains abundant evidence of the immune status following infection. Cytokines, chemokines, and antibodies can be assayed to determine the extent to which a patient responded to a pathogen. We examined serum and plasma from a cohort of patients infected with SARS-CoV-2 early in the pandemic and compared them to negative-control sera. Cytokine and chemokine concentrations varied depending on the severity of infection, and antibody responses were significantly increased in severe cases compared to mild to moderate infections. Neutralization data revealed that patients with high titers against an early 2020 isolate had detectable but limited neutralizing antibodies against newly circulating SARS-CoV-2 variants of concern. This study highlights the potential of re-infection for recovered COVID-19 patients.

## Introduction

In December 2019, severe acute respiratory syndrome coronavirus 2 (SARS-CoV-2) was first reported in the city of Wuhan, Hubei province, China, causing variably severe respiratory tract pathology termed coronavirus disease 2019 (COVID-19). COVID-19 is often a mild disease associated with low-grade fever and loss of taste and smell. However, critical cases of COVID-19 do occur, and are characterized by severe pneumonia and acute respiratory distress syndrome (1) leading to organ failure and death (2). As of October 20^th^ 2021, over 241 million cases have been reported worldwide, and over 4.9 million people have died of COVID-19 (https://coronavirus.jhu.edu/map.html).

The spectrum of disease caused by SARS-CoV-2 ranges from no or mild to critical. Mild to moderate cases are characterized by mild symptoms ranging to mild pneumonia and account for up to 81% of infections. Severe cases account for 14% of cases, which involve dyspnea, hypoxia, or greater than 50% lung involvement as determined by imaging. Five percent of patients are deemed critical based on conditions of respiratory failure, shock, or multiorgan system dysfunction (3, 4). In many severely affected patients, SARS-CoV-2 infection triggers an overactive immune response known as a “cytokine storm.” Immune cells produce high levels of inflammatory cytokines leading to systemic shock and death (5). As such, cytokines have been studied extensively in the context of SARS-CoV-2 infection and have been found to be central to the pathophysiology of COVID-19 (6, 7).

A thorough understanding of appropriate immune responses is vital to the development of effective medical intervention strategies and vaccines. Besides cytokine and chemokine production following infection, antibodies generated by COVID-19 patients have been studied and reported in detail. Infection with SARS-CoV-2 has been found to induce non-class-switched, class-switched, and neutralizing antibodies in immunocompetent patients (8-12). The long term stability of the antigen-specific and neutralizing antibody response has been found to be up to 13 months in patients (13-16). Pre-existing antibody populations may also contribute to disease severity such as autoantibodies to type I interferons (17). As SARS-CoV-2 mutates, changes to the sensitivity of pre-exisitng neutralizing antibody populations may be effected (18). As such, the beta and delta variants both have displayed decreased sensitivity to pre-existing neutralizing antibodies (15, 19-21).

In this study, we evaluated 131 serum and plasma samples from 55 COVID-19 patients alongside serum and plasma from 20 uninfected patients for the presence of 38 cytokines and chemokines, anti-SARS-CoV-2 spike protein-specific IgG, and neutralizing antibodies. Our results indicate that infection with SARS-CoV-2 results in changes in a number of cytokines and chemokines that correlate to disease severity. We also found that COVID-19 patients exhibit increased titers of antigen-specific IgG and neutralizing antibody titers compared to uninfected individuals. Furthermore, we determined that the neutralizing activity of our sample cohort extended to three new SARS-CoV-2 variants of concern (VOC), Alpha (α; B.1.1.7), Beta (β; B.1.351), and Delta (δ; B.1.617.2) which emerged months after the start of the pandemic. This study corroborates previous data examining serum concentrations of cytokines, chemokines, and antigen-specific antibodies in COVID-19 patients. Most importantly, it highlights the cross-reactive neutralization capabilities of unvaccinated COVID-19 survivors against emerging SARS-CoV-2 variants and the potential for re-infection.

## Materials and Methods

### Cells and Viruses

Vero E6 cells (mycoplasma negative) were grown at 37°C in 5% CO_2_ in Dulbecco’s modified Eagle’s medium (DMEM) (Sigma-Aldrich, St. Louis, MO) containing 10% fetal bovine serum (FBS) (Wisent Inc.), 2 mM L-glutamine (Thermo Fisher Scientific, Waltham, MA), 50 U/mL penicillin (Thermo Fisher Scientific), and 50 μg/mL streptomycin (Thermo Fisher Scientific). SARS-CoV-2 isolate nCoV-WA1-2020 (MN985325.1) (22), SARS-CoV-2 isolate B.1.351 (hCoV-19/South African/KRISP-K005325/2020), SARS-CoV-2 isolate B.1.1.7 (hCOV_19/England/204820464/2020), and SARS-CoV-2 isolate B.1.617.2 (hCoV-19/USA/KY-CDC-2-4242084/2021) were used for the neutralizing antibody assays. The following reagent was obtained through BEI Resources, NIAID, NIH: Severe Acute Respiratory Syndrome-Related Coronavirus 2, Isolate hCoV-19/England/204820464/20200, NR-54000, contributed by Bassam Hallis. SARS-CoV-2 B.1.351 was obtained with contributions from Dr. Tulio de Oliveira and Dr. Alex Sigal (Nelson R Mandela School of Medicine, UKZN). SARS-CoV-2 B.1.617.2 was obtained with contributions from B. Zhou, N. Thornburg and S. Tong (Centers for Disease Control and Prevention, USA). All viruses were grown and titered on Vero E6 cells, and sequence confirmed.

### Serum and plasma samples

A total of 131 serum and plasma samples collected from 75 unique individuals were analyzed in this study. All samples were either remnant sera or plasma (from EDTA-anticoagulated whole blood), originally collected for standard-of-care diagnostic testing from inpatients being treated for COVID-19 (n = 55 [73%]) or were from SARS-CoV-2-uninfected volunteers (n = 20 [27%]; referred to as “normal” samples). Of the patients, the average age was 58 years (range 13-93 years), 25 (45%) were female and 27 (49%) had more than one specimen assessed. Based on review of clinical charts and according to CDC criteria (www.cdc.gov/coronavirus/2019-ncov), patients were grouped into three illness severity categories: Mild to moderate (mild symptoms to mild pneumonia), severe (dyspnea, hypoxia or more than 50% lung involvement on imaging), and critical (respiratory failure, shock or multiorgan system dysfunction). Of the normal samples, volunteers who provided a one-time sample were an average age of 44 years (range 26-89 years) and 15 (75%) were female. All samples, including patients and volunteers, were deidentified and assigned study-specific identifiers to protect patient confidentiality. Samples were then aliquoted and frozen at −80°C until shipment to the Rocky Mountain Laboratories for analysis. Samples were γ-irradiated (4 MRad) to inactivate potential infectious pathogens upon receipt and prior to analysis. This work was approved by the Indiana University Institutional Review Board (IRB# 2004155084).

### *Cytokine* analysis

Samples were diluted 1:2 in serum matrix for analysis with Milliplex Human Cytokine/Chemokine Magnetic Bead Panel as per manufacturer’s instructions (EMD Millipore Corporation). Concentrations for analytes (EGF, FGF-2, Eotaxin, TGF-α, G-CSF, Flt-3L, GM-CSF, Fractalkine, IFNα2, IFNγ, GRO, IL-10, MCP-3, IL-12p40, MDC, IL-12p70, IL-13, IL-15, sCD40L, IL-17A, IL-1RA, IL-1α, IL-9, IL-1β, IL-2, IL-3, IL-4, IL-5, IL-6, IL-7, IL-8, IP-10, MCP-1, MIP-1α, MIP-1β, TNFα, TNFβ, and VEGF) were determined for all samples using the Bio-Plex 200 system (BioRad Laboratories, Inc.).

### Antibody level determination

Antibody titers were determined using enzyme-linked immunosorbent assay (ELISA). Flat-bottom immuno 96-well plates (Nunc Maxisorp, Thermo Fisher Scientific) were coated overnight with 1 ug/ml SARS-CoV-2 (2019-nCoV) Spike Receptor Binding Domain (polyhistidine-tagged) recombinant protein (Sino Biological) diluted in PBS. Plates were washed and blocked the following day with 3% milk. After washing, serum and plasma samples were diluted 1:100, and then serially diluted 1:4 in 1% milk and incubated for one hour at room temperature. Plates were washed before addition of peroxidase-labeled anti-human IgG (KPL). Following a one-hour incubation at room temperature, plates were washed and ABTS 2-component Microwell Peroxidase Substrate (SeraCare) was added. Plates were incubated for 30 minutes in the dark before being read at 405 nm on a GloMax Explorer (Promega).

### *Neutralizing antibody* assay

Neutralization antibody assays were performed as detailed in van Doremalen et al. (1). Briefly, serum and plasma samples were heat-inactivated for 30 minutes at 56°C. They were diluted 1:10 and then 1:2 for subsequent dilutions. SARS-CoV-2 virus stocks were diluted to 2,000 TCID50/ml and 70 μl was then added to each well of diluted sample. Following a one-hour incubation at 37°C, the serum-virus mixture was transferred to 96-well plates containing high-passage Vero E6 cells. After six days, cytopathic effect (CPE) was read. The virus neutralization titer was determined to be the lowest concentration of serum antibody where CPE was not observed.

### *Statistical* analysis

Statistical analysis was performed using Prism 8. Statistically significant differences between groups for cytokines were determined using one-way ANOVA; IgG and neutralzing titers were evaluated applying Mann-Whitney test. Significance is indicated as follows: p<0.0001 (****), p<0.001 (***), p<0.01 (**) and p<0.05 (*).

## Results

### COVID-19 patients exhibit different levels of cytokines and chemokines, which correlate with disease severity

We received 111 patient serum and plasma samples that were categorized according to CDC guidelines into mild to moderate and critical cases. In addition, we obtained 20 serum and plasma samples from healthy adult volunteers designated “normal” controls in our studies. We first sought to determine the circulating immune status by assessing the presence of 38 different cytokines and chemokines in the serum and plasma of patients infected with SARS-CoV-2, alongside uninfected volunteers. We found that infection with SARS-CoV-2 resulted in significant changes in multiple cytokines and chemokines compared to negative control serum and plasma (Figure 1). This phenomenon was evident in both mild to moderate and critical infections. For instance, serum and plasma from patients with a mild to moderate infection contained significantly greater levels of MCP-3, IL-1α, TNFβ, IL-4, IL-5, IL-6, IL-8, IL-9, and IL-13 compared to serum and plasma from patients that were critically ill (Figure 1A). Mild to moderate infections also showed significant increases in these cytokines, along with IL-15, compared to healthy adults (Figure 1B). Critical infections resulted in significantly increased levels of sCD40L, IP-10, and IL-15 compared to normal controls (Figure 1 B). All of the other cytokines and chemokines tested showed no significant differences between control and infected patients (Supplementary Figure 1).

**Figure 1.**
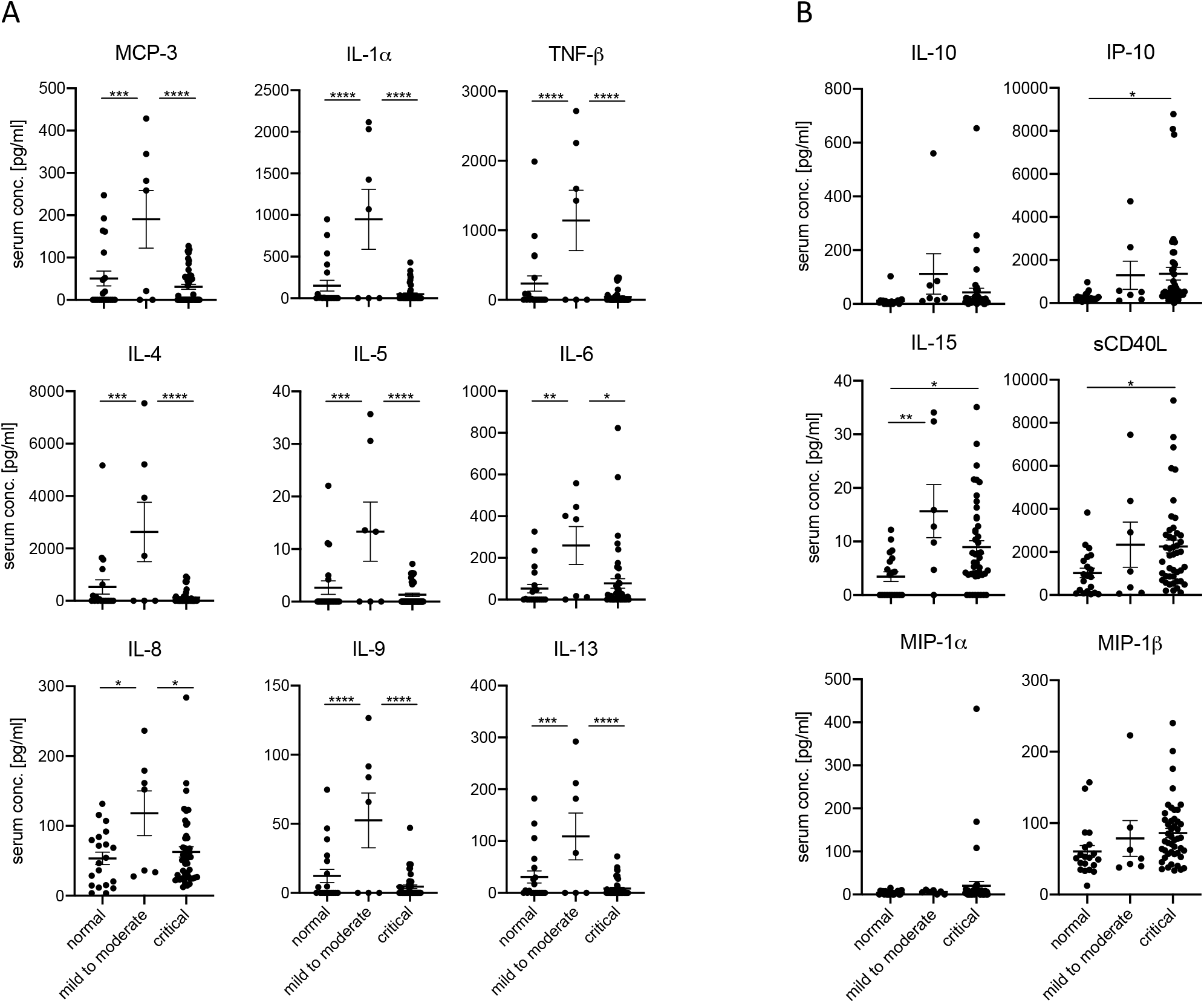
COVID-19 patients exhibit varying levels of cytokines and chemokines that correlate with disease severity. Cytokine and chemokine levels for day-of-admission COVID-19 serum samples alongside 20 normal samples. Error bars represent standard error. Statistically significant differences as determined by one-way ANOVA are indicated as p<0.0001 (****), p<0.001 (***), p<0.01 (**), and p<0.05 (*).

### Cytokine levels remain steady when measured over time

Nine of the COVID-19-infected patients were sampled more than once. We determined longitudinally whether cytokine and chemokine concentrations varied over time. Overall, there were no significant changes in any of the evaluated cytokines or chemokines during the periods of time that were sampled (Supplementary Figure 2).

### Failure to recover from critical COVID-19 is correlated with increased levels of IL-6 and IP-10 coupled with insufficient levels of sCD40L

Amongst critical patients, we found that the serum and plasma of those who succumbed to the infection contained significantly more IL-6 and IP-10 than control patients. Additionally, patients who survived infection had significantly increased sCD40L compared to normal controls (Figure 2). Although serum and plasma from fatal disease patients contained more sCD40L than serum and plasma of normal controls, it was not significantly increased relative to serum and plasma from patients who recovered from infection, indicating that this soluble mediator may be important for surviving COVID-19.

**Figure 2.**
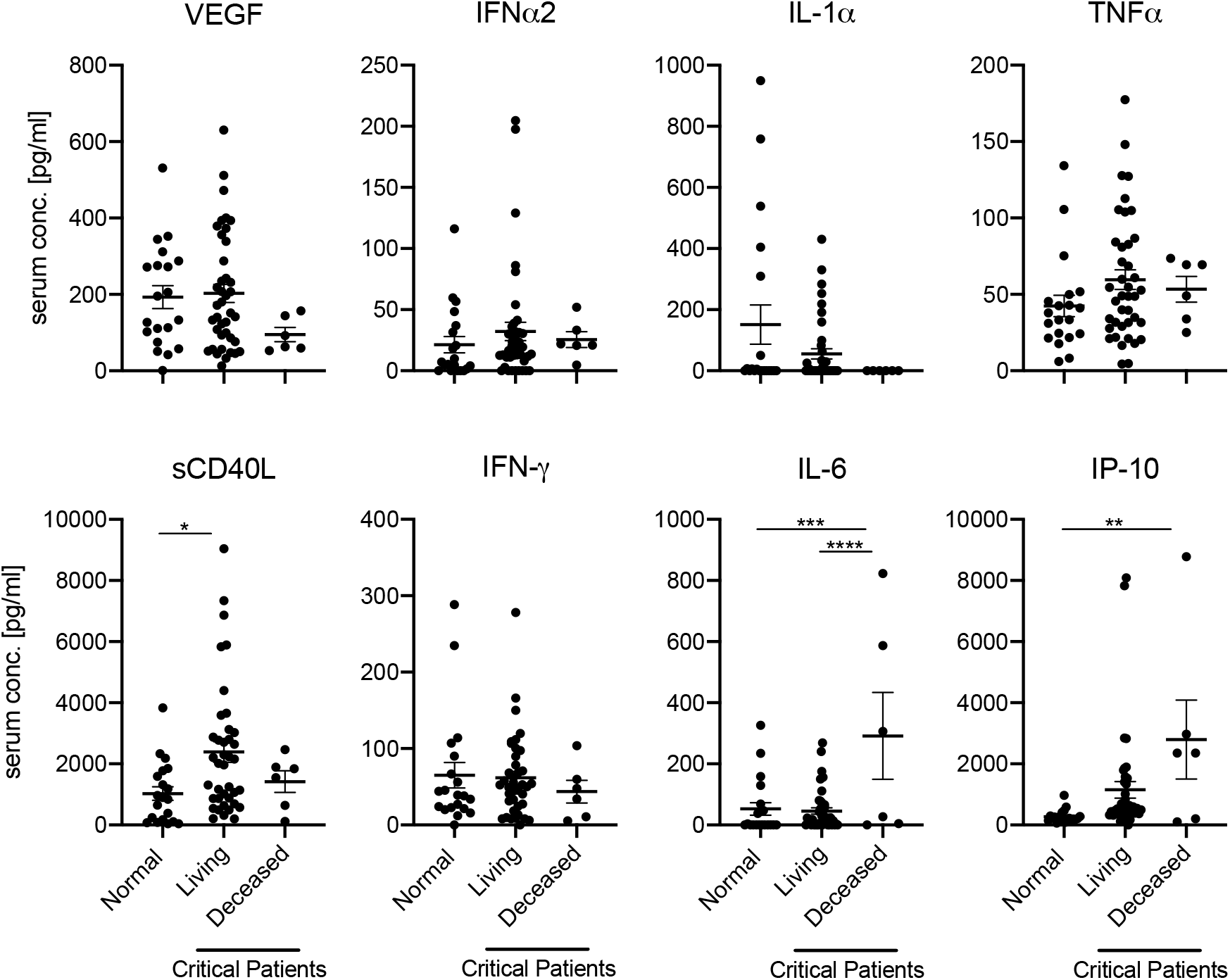
Serum IL-6, IP-10, and sCD40L predict disease outcome in COVID-19 patients. Cytokine and chemokine levels for day-of-admission critically infected COVID-19 patients who either recovered from infection (living) or failed to recover (deceased), alongside 20 uninfected (normal) patients. Error bars represent standard error. Statistically significant differences as determined by one-way ANOVA are indicated as p<0.0001 (****), p<0.001 (***), p<0.01 (**), and p<0.05 (*).

### *Critically infected patients develop strong anti-SARS-CoV-2 IgG and neutralizing antibody* responses

Next, we assessed whether serum and plasma of COVID-19 patients contained IgG specific for the receptor-binding domain (RBD) of the SARS-CoV-2 spike protein. We found that half of the patients who had a mild to moderate infection produced RBD-specific IgG, and the data was statistically significant compared to healthy controls (Figure 3A). In contrast, critically-infected patients developed high titers of RBD-specific IgG, which was significantly greater than both healthy controls and those with mild to moderate COVID-19. The presence of RBD-specific IgG did not predict disease outcome, as there was no significant difference in titers between patients that recovered from infection and those that did not (Figure 3B).

**Figure 3.**
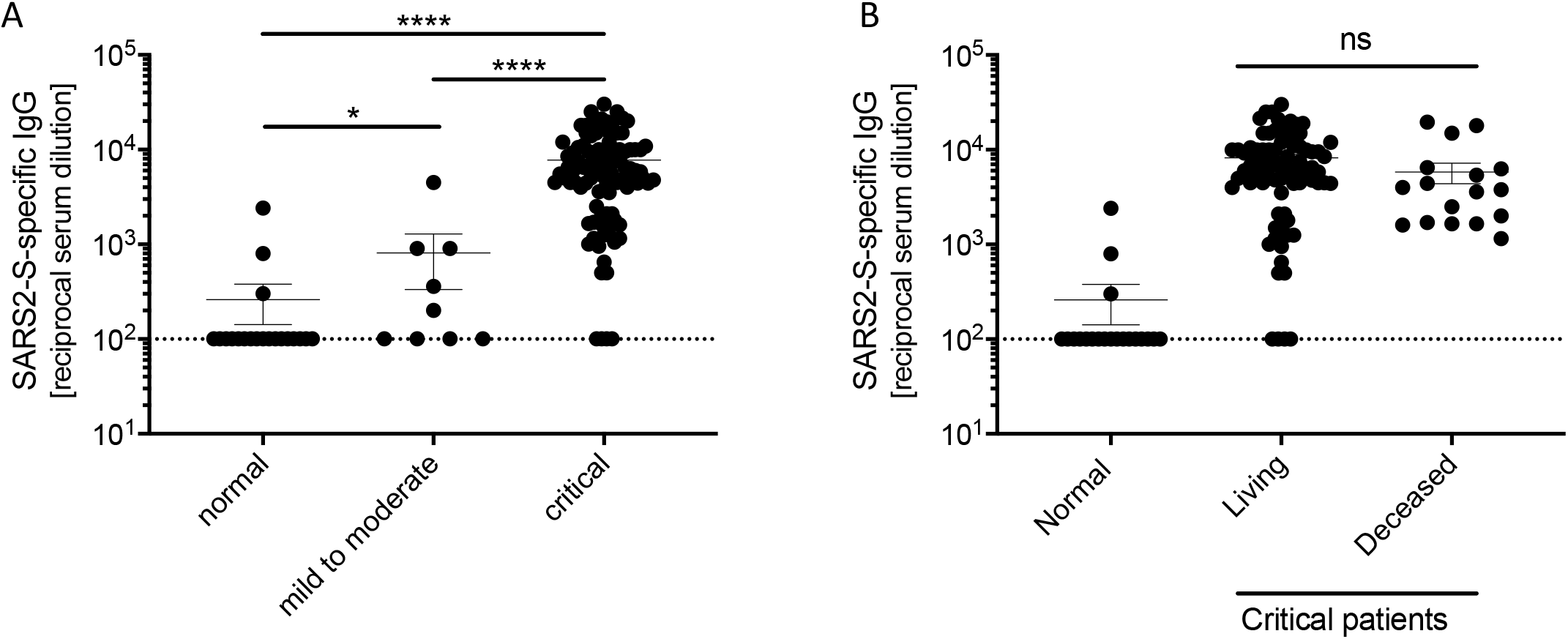
Critical COVID-19 induces high titers of anti-SARS-CoV-2 IgG. **(A)** IgG titers specific to SARS-CoV-2 spike receptor-binding domain for COVID-19 and normal serum samples. **(B)** IgG titers of critically infected COVID-19 patients who either recovered from infection (living) or failed to recover (deceased), alongside 20 uninfected (normal) patients. Error bars represent standard error. Statistically significant differences as determined by Mann-Whitney test are indicated as p<0.0001 (****), p<0.001 (***), and p<0.05 (*).

Three of the healthy control serum and plasma samples contained SARS-CoV-2-specific IgG without COVID-19 medical history. We postulated that the presence of IgG may not translate to the ability to neutralize SARS-CoV-2. Therefore, we assessed the serum and plasma for neutralizing antibodies against SARS-CoV-2. The majority of control serum and plasma did not contain detectable levels of neutralizing antibodies, with the exception of one patient who exhibited a detectable, albeit very low, titer (Figure 4A). However, mild to moderate infection led to the production of significantly higher levels of neutralizing antibody titers compared to controls. Patients that were critically infected exhibited high titers of neutralizing antibodies, approximately two logs greater than healthy controls and one-and-a-half logs higher than patients with mild to moderate disease.

**Figure 4.**
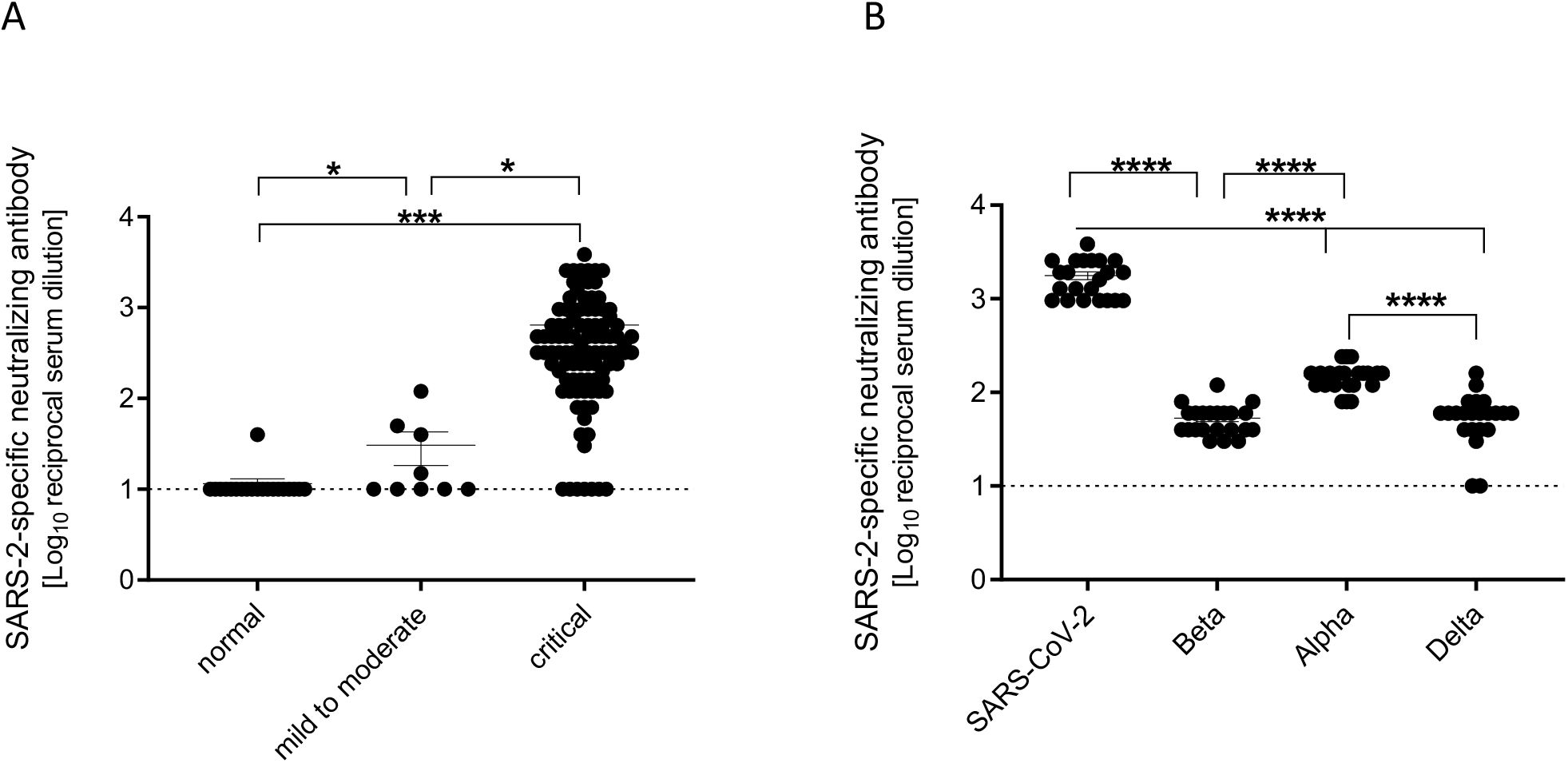
Critical COVID-19 induces high titers of anti-SARS-CoV-2 neutralizing antibodies. **(A)** SARS-CoV-2-specific neutralizing antibody titers for COVID-19 and normal serum samples. **(B)** Neutralizing antibody titers for COVID-19 serum samples comparing SARS-CoV-2 variants USA-WA1, Beta, Alpha, and Delta. Error bars represent standard error. Statistically significant differences as determined by Mann-Whitney test are indicated as p<0.0001 (****), p<0.001 (***), and p<0.05 (*).

Some COVID-19 patients that participated in this study were sampled repeatedly over the course of infection and recovery. Therefore, we determined whether their neutralizing antibody titers remained stable over time. We found that these patients continued to produce neutralizing antibodies against SARS-CoV-2 over the course of the disease (Supplementary Figure 3). Antibody titers remained nearly the same, exhibiting less than a log change, from the first to the final day of sampling (up to nine days post-admission). Unfortunately, no further time points were sampled from the patients.

### COVID-19 serum and plasma contains neutralizing antibodies against VOC

Finally, we sought to determine whether human COVID-19 serum and plasma contains antibodies capable of neutralizing three VOC that emerged later in the pandemic – Alpha, Beta, and Delta variants. Therefore, we tested serum and plasma samples that contained high titers of neutralizing antibodies against the original virus, but this time we assessed neutralizing antibodies against the 3 VOC. We found that the serum and plasma contained antibodies capable of neutralizing these SARS-CoV-2 variants, but the antibody titers were significantly lower than the titers against the original virus (Figure 4B). Interestingly, the titers against the Alpha variant were significantly higher than those for the Beta and Delta variants indicating limited protection from re-infection with the currently circulating Delta variant (Figure 4B).

## Discussion

Since the beginning of the COVID-19 pandemic, clinicians and scientists have sought to investigate the components of patient serum for evidence of either sufficient or aberrant immune responses. Components of serum have been shown to be effective in the treatment of those suffering from COVID-19, as convalescent plasma infusion can lead to a decrease in the severity of disease (23, 24). Understanding the difference between an immune response that leads to recovery from infection and one that leads to a negative outcome (aberrant) is essential in the design of treatments and vaccines, which are necessary to bring an end to this devastating pandemic.

One avenue taken by investigators has been to explore the presence or absence of various cytokines and chemokines in patient serum. Severe disease has been associated with an aberrant immune response termed “cytokine storm,” which is characterized by an overactivation of the immune system leading to exaggerated levels of cytokines released into the circulation. Multiorgan dysfunction and failure associated with septic shock can be fatal (5, 25). In contrast, those with mild disease exhibit functional immune responses characterized by appropriate levels and types of cytokines, leading to disease resolution (7). One cytokine that has been highlighted amongst research studies is IL-6. For instance, Herold et al. found that IL-6 was a key predictor of respiratory failure in hospitalized COVID-19 patients (26). Other studies have yielded similar results, indicating a role for IL-6 in the severity and outcome of the disease (27-29). Our study supports previous studies showing that high levels of IL-6 lead to poor outcomes for COVID-19 patients.

Interestingly, our data show that moderate levels of key cytokines and chemokines are evident in mild to moderate cases of COVID-19. It is the “Goldilocks” phenomenon: too much or too little of some cytokines is not good; rather, the levels must be “just right”. In support of this concept, Yang et al. observed that serum IL-1β, IL-1Ra, IL-6, IL-7, IL-10, IP-10, and TNF-α are all important in classifying COVID-19 cases into mild, moderate, and severe (30). This study also found that IP-10 was significantly higher in severe cases of COVID-19 compared to mild cases. Another study found that IP-10 levels were highest in patients that required ICU admission (31). Our study supports the previous work finding very high levels of IP-10 in the serum of patients who succumbed to SARS-CoV-2 disease. Serum antibodies are known to be important for both protection and treatment of COVID-19. Effective humoral immune responses to vaccination or infection lead to the production of neutralizing antibodies that contribute to clearance of the virus. Our data demonstrate that infection with SARS-CoV-2 results in increased levels of antigen-specific IgG, and that severe infection leads to higher levels compared to mild to moderate infection, corroborating other studies. For instance, Chen et al. determined that symptom severity correlated directly with the magnitude and durability of class-switched serum antibodies, as well as other studies (13, 32, 33). However, little is known if the magnitude of antibody level correlates with re-infection potential particularly with VOC.

Although presence of IgG is evidence of an effective immune response, it is important to decipher whether these antibodies are capable of neutralizing virus. It has been found that not all recovered COVID-19 patients develop sufficient neutralizing antibody titers (12). Our study showed that while SARS-CoV-2 cross-reactive IgG antibodies were present in control patients, these antibodies were not capable of neutralizing SARS-CoV-2. A recent study suggests that exposure to a seasonal coronavirus can induce the production of antibodies against SARS-CoV-2, but these antibodies are not protective against the virus (34). Therefore, detection of IgG alone cannot always predict protection from reinfection. This is an important distinction that clinicians need to make when examining data from recovered patients. Optimally, a test to determine the presence of specific neutralizing antibodies would be more informative than our current ELISA, which only detects antibodies that are specific for SARS-CoV-2.

A key feature of SARS-CoV-2 is its high mutation rate (35, 36). Selective pressures acting on the virus have led to mutations that allow the virus to spread more efficiently and to evade host immune responses (37). Fortunately, we and others have found that there appears to be some albeit limited cross-protection against the circulating VOC (38, 39). In our hands, patients who survived infection with the original SARS-CoV-2 generate antibody responses capable of neutralizing three different VOC, and these antibodies are more effective against the Alpha variant compared to the Beta and Delta variants. It is important to note that the cross-reactive neutralizing potential from the original SARS-CoV-2 was significantly less for all three of the VOC tested. However, recent studies provide hope that even low levels of neutralizing antibodies will lead to better outcomes after re-infection with VOC for patients who have been vaccinated or survived natural infection with SARS-CoV-2 early in the pandemic (38, 40, 41). It has been demonstrated that individuals whom have been previously infected more rapidly develop neutralizing antibodies post-vaccination with an mRNA vaccine. The neutralizing antibody titer is blunted across multiple VOC with the Beta and Gamma variants have the most dramatic decrease followed by the Delta variant (42). The decrease in neutralization efficiency can be attributed to the mutations the spike protein has acquired, specifically the dominant epitopes that are targeted by neutralizing antibodies to each variant. It has recently been shown that patients infected with an earlier isolate, similar to the ancestral WA1 isolate we used in this study, develop neutralizing antibodies against class 2 epitopes, while patients infected with the Beta variant develop neutralizing antibodies against class 3 epitopes (43). It would be interesting to know if protection against newly emerging VOC is enhanced by the neutralizing antibody response in one group or the other.

Our study provides additional support for the growing body of literature examining human COVID-19 serum samples. Our data supports established work that increased levels of IL-6 and IP-10 contribute to enhanced disease phenotype. In addition, our study highlights the importance of both the antigen-specific antibody response and its functionality to neutralize emerging VOC. The more fully we understand effective immune responses to this pathogen, the greater our ability to successfully treat those who are infected, and vaccinate those we hope to protect against infection.

## Acknowledgements

We thank members of the Molecular Pathogenesis Unit, Virus Ecology Section, and Research Technology Branch (all NIAID) for their efforts to obtain and characterize the SARS-CoV-2 isolates.

## Author contributions

A.M. and R.F.R. conceived the idea. A.M. secured funding. A.J.G., R.F.R., and A.M. designed the experiments. P.M.R., J.-P.L., M.K.Z., and R.F.R. collected and provided the serum samples and cooresponding clinical chart data. A.J.G., and K.L.O conducted the experiments and acquired the data. A.J.G., K.L.O, K.S., R.F.R. and A.M. analyzed and interpreted the data. A.J.G., K.L.O., and A.M. prepared the manuscript. All authors approved the submitted manuscript.

## Data availability

All data is available in the manuscript.

---

**Supplementary Figure 1.**
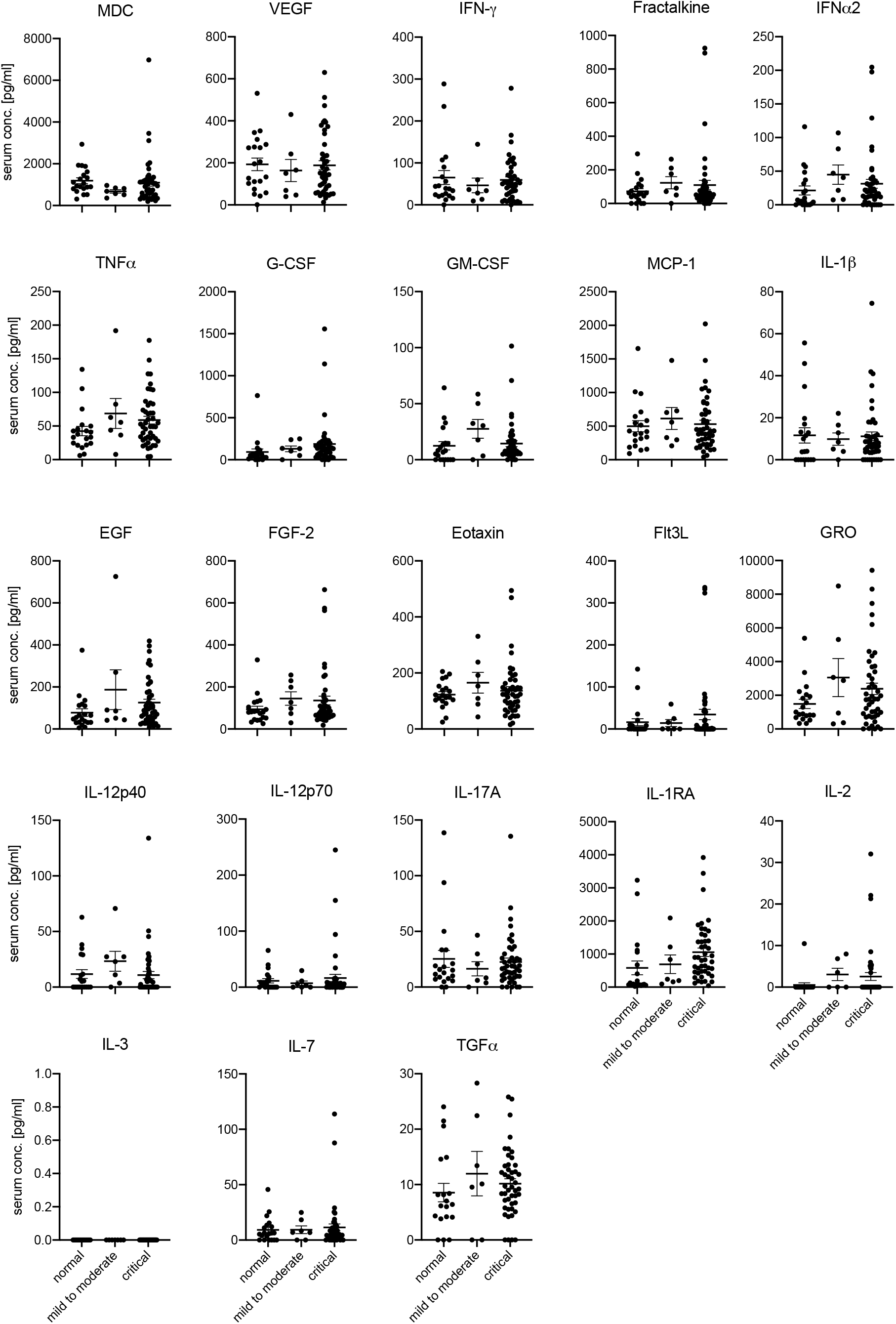
Cytokine and chemokine levels for day-of-admission COVID-19 serum samples alongside 20 normal samples. Error bars represent standard error.

**Supplementary Figure 2.**
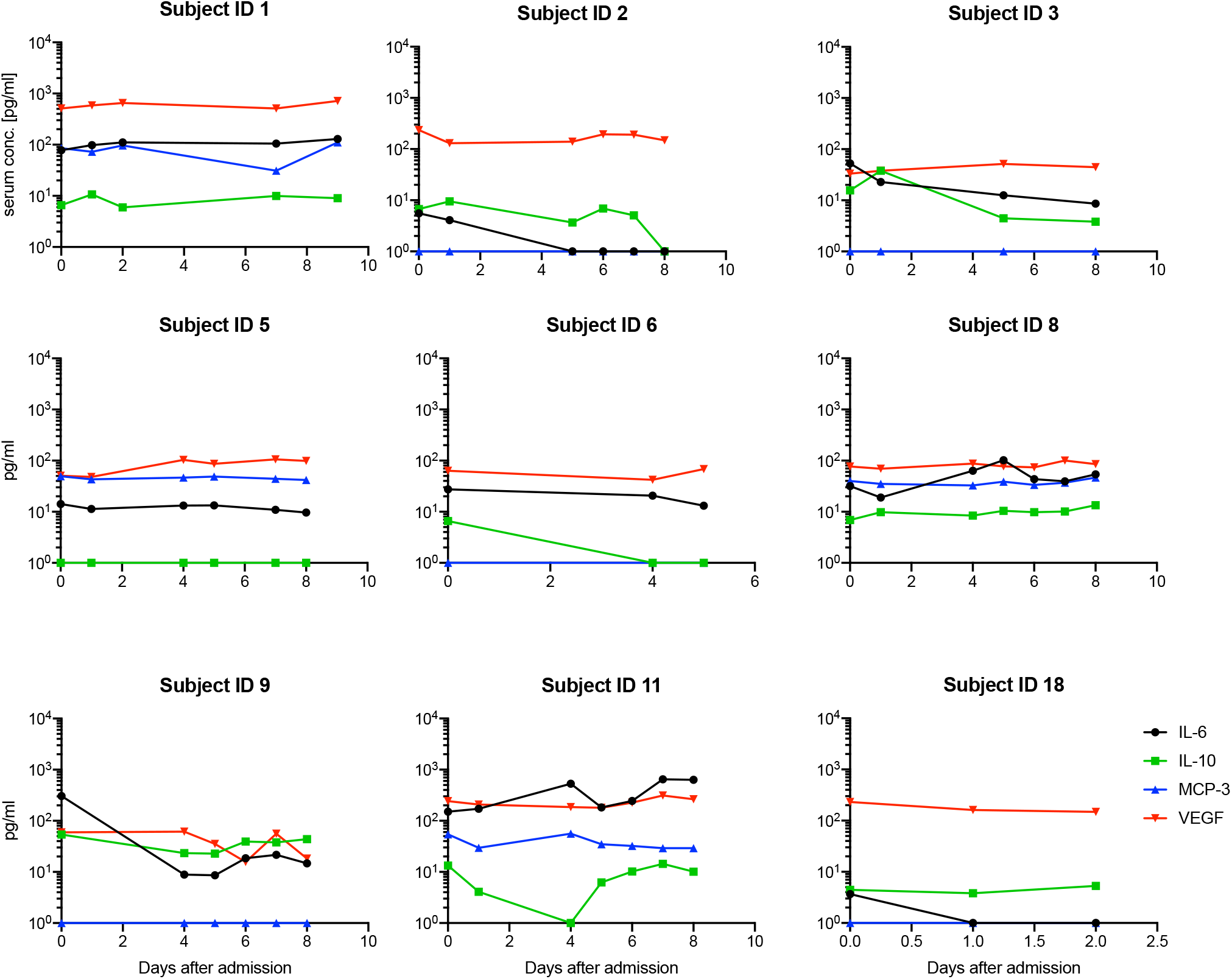
Cytokine and chemokine levels of selected patients over time.

**Supplementary Figure 3.**
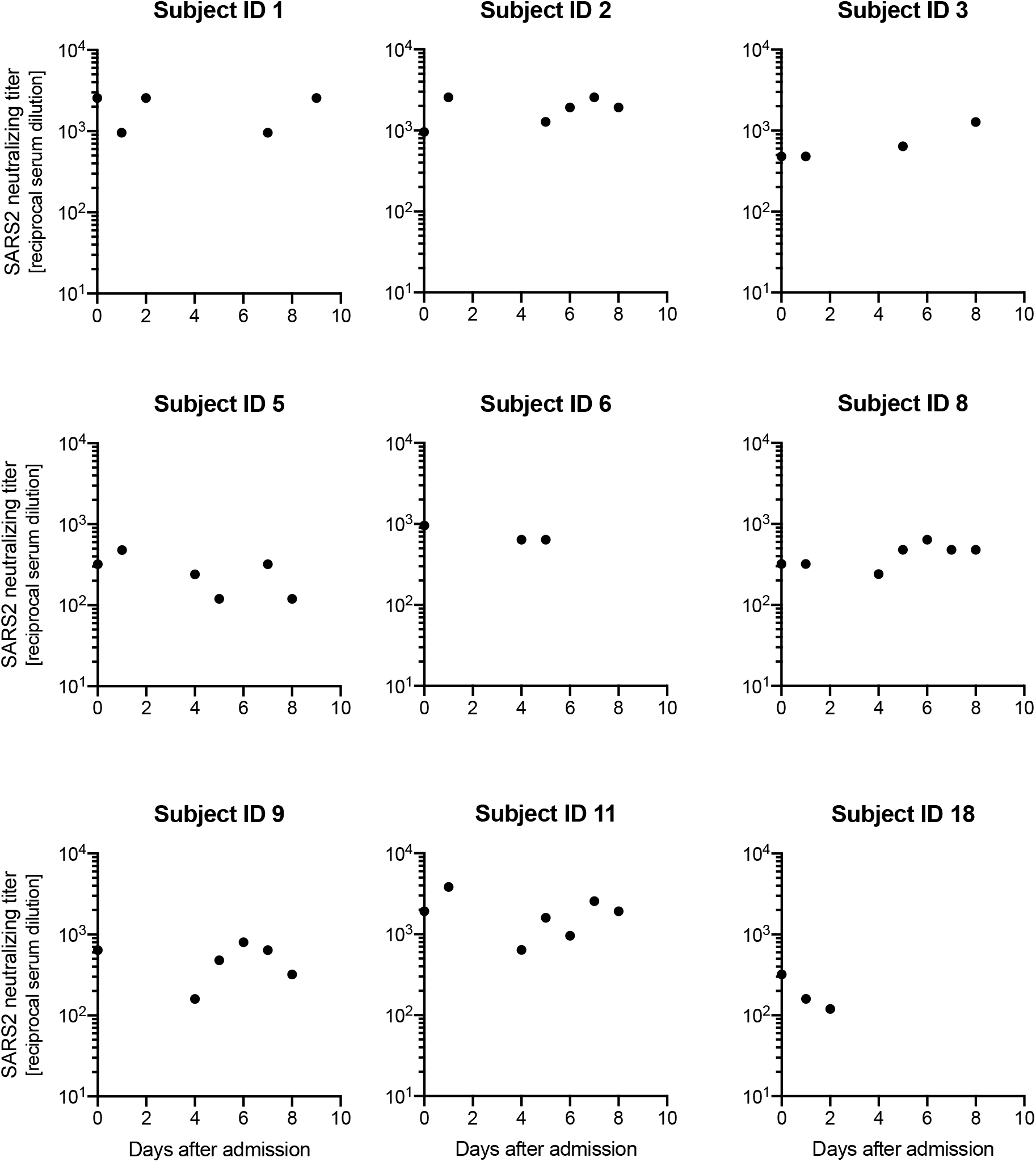
Anti-SARS-CoV-2 neutralizing antibodies remain steady over time in selected patients.

